# Functional implications of impaired dynamic cerebral autoregulation in young healthy women; a comparative investigation

**DOI:** 10.1101/406827

**Authors:** Lawrence Labrecque, Kevan Rahimaly, Sarah Imhoff, Myriam Paquette, Olivier Le Blanc, Simon Malenfant, Audrey Drapeau, Jonathan D. Smirl, Damian M. Bailey, Patrice Brassard

**Affiliations:** Department of Kinesiology, Faculty of Medicine, Université Laval, Québec, Canada; Research center of the Institut universitaire de cardiologie et de pneumologie de Québec, Québec, Canada; Concussion Research Laboratory, Health and Exercise Sciences, University of British Columbia Okanagan, British Columbia, Canada; Neurovascular Research Laboratory, Faculty of Life Sciences and Education, University of South Wales, United Kingdom

**Keywords:** brain, sexual differences, cerebral blood flow, dynamic cerebral

## Abstract

Women exhibit higher prevalence of orthostatic hypotension with presyncopal symptoms compared to men. These symptoms could be influenced by an attenuated ability of the cerebrovasculature to respond to rapid changes in blood pressure (BP) [dynamic cerebral autoregulation (dCA)]. However, the influence of sex on dCA remains equivocal. We compared dCA in 11 women (25 ± 2 y) and 11 age-matched men (24 ± 1 y) using a multimodal approach including a sit-to-stand maneuver and forced oscillations (5 min of squat-stand performed at 0.05 and 0.10 Hz). The prevalence of initial orthostatic hypotension (IOH; decrease in systolic ≥ 40 mmHg and/or diastolic BP ≥ 20 mmHg) during the first 15 sec of sit-to-stand was determined as a functional outcome. In women, the decrease in mean middle cerebral artery blood velocity (MCAv_mean_) following the sit-to-stand was greater (−20 ± 8 vs. -11 ± 7 cm sec^-1^; *p=0.018*) and the onset of the regulatory change (time lapse between the beginning of the sit-to-stand and the increase in the conductance index (MCAv_mean_/mean arterial pressure(MAP)) was delayed *(p=0.007).* Transfer function analysis gain during 0.05 Hz squat-stand was ∼48% higher in women (6.4 ± 1.3 vs. 3.8 ± 2.3 sec; *p=0.017*). The prevalence of IOH was comparable between groups (4/9 vs. 5/9, *p=0.637).* These results indicate the cerebrovasculature of healthy women has an attenuated ability to react to large and rapid changes in BP in the face of preserved orthostasis, which could be related to a higher cerebrovascular reserve to face a rapid transient hypotension.

**NEWS & NOTEWORTHY:** The novel findings of this study are that healthy women have impaired dynamic cerebral autoregulation, although the prevalence of orthostatic intolerance was similar in women and men. These results indicate the cerebrovasculature of healthy women has an attenuated ability to react to large and rapid changes in blood pressure in the face of preserved orthostasis, which could be related to a higher cerebrovascular reserve to face a rapid transient hypotension.

## INTRODUCTION

The prevalence of orthostatic hypotension is higher in women compared to men (13). In addition, women suffer more often from symptoms of cerebral hypoperfusion such as light-headedness, nausea and blurred vision (3). These symptoms could be influenced by an attenuated ability of the cerebrovasculature to respond to rapid changes in arterial blood pressure [traditionally referred to as dynamic cerebral autoregulation (dCA)].

Accumulating evidence supports the notion that cerebral blood flow (CBF) is regulated differently in women compared to men. Resting CBF (21) and cerebrovascular reactivity to carbon dioxide (15) are higher in women. Although some disparities seem to exist in regards to CA (35), very few studies have attempted to assess this crucial CBF determinant in healthy young women. Using transfer function analysis (TFA) of spontaneous or forced oscillations in mean arterial pressure (MAP) and middle cerebral artery blood velocity (MCAv), investigators reported older women have either similar (25) or enhanced dCA (9, 11, 37) compared to age-matched men. Nevertheless, these findings are difficult to translate to a younger population since studies have shown older women regulate CBF in a different manner when compared with younger women (11).

Most metrics quantifying dCA have a limited ability to characterize each other (31). Furthermore, characterization of dCA employing diverse analytical techniques can produce variable physiological interpretations (31, 32). Thus, when performing investigations of dCA, the utilization of a multimetrics approach could help improve our understanding of this response. Previously, we have employed this approach to study the influence of cardiorespiratory fitness (CRF) on dCA in healthy fit men (18). This approach revealed CRF is associated with an intact ability of the cerebrovasculature to dampen spontaneous oscillations in MAP (comparable TFA metrics between fit men vs. controls). This approach also included forced BP oscillations using repeated squat-stand maneuvers, to improve the interpretation of the linear association between BP and MCAv (28), which revealed a reduced capability of reacting to large and rapid changes in MAP (delayed onset of the regulatory response; increased absolute TFA gain during 0.10 Hz repeated squat-stand maneuvers). These results highlight the importance of including BP stimuli of different natures and magnitudes when examining the capability of the cerebrovasculature to respond to changes in MAP (27).

Therefore, the aim of this study was to examine to what extent sex potentially influences dCA in a young and healthy population using a multiple assessment strategy and hemodynamic stressors (sit-to-stand and TFA of forced MAP and MCAv oscillations). We also determined the prevalence of initial orthostatic hypotension (IOH) as a functional outcome, in order to appreciate how the potential impact of sex on dCA translates in terms of physiological outcome. We hypothesized women would have better dCA compared to men, and dCA metrics would be related to orthostatic tolerance.

## MATERIALS AND METHODS

### Ethics and informed consent

All participants provided informed consent prior to participating in the investigation, and the study was approved by the Comité d’éthique de la recherche de I’lUCPQ-Université Laval (CER: 20869 and 21180).

### Participants

Twenty-two moderately-trained endurance athletes were recruited for this study: eleven women [maximal oxygen consumption 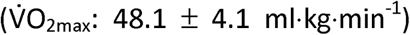] and eleven men 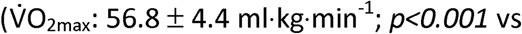. Women and men were matched for age, body mass index (BMI) and volume of weekly training. All the participants competed in a variety of endurance-based sports including cycling (women: n=1; men: n=5), triathlon (women: n=4; men: n=5), mountain biking (women: n= 1), running (women: n= 4) and cross-country skiing (women: n=1; men: n=1). All participants were free from any medical conditions, demonstrated a normal 12-lead ECG, and were not taking any medications. Two women were taking oral contraceptive continuously since > 1 year and two women had an intrauterine device. The remaining women were tested during menses or the early follicular phase (days 1 to 10) of their menstrual cycle (n=7).

### Experimental protocol

Parts of this experimental protocol, including dCA and orthostatic tolerance metrics from the group of eleven men included in this analysis have previously been published (18), as part of an investigation with a separate focus on the influence of CRF on dCA. Although the current study employed a similar experimental design, it represents a separate question (influence of sex on dCA) through the addition of a group of age and BMI matched women. As previously described (18), participants visited the laboratory on two occasions to perform: 1) an incremental cycling test for 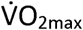 determination, and 2) anthropometrics, resting measurements and the evaluation of dCA and orthostatic tolerance. Participants were asked to avoid exercise training for at least 12 h, as well as alcohol and caffeine consumption for 24 h before each visit. All sessions and evaluations were executed in the exact same order for all participants and there was at least 48 h between testing sessions.

### Measurements

#### Systemic hemodynamics

Heart rate (HR) was measured using a 5-lead ECG. Beat-to-beat BP was measured by the volume-clamp method using a finger cuff (Nexfin, Edwards Lifesciences, Ontario, Canada). The cuff was placed on the right middle finger and referenced to the level of the heart using a height correct unit for BP correction. MAP was obtained by integration of the pressure curve divided by the duration of the cardiac cycle. This method has been shown to reliably index the dynamic changes in beat-to-beat BP which correlate well with the intra-arterial BP recordings and can be used to describe the dynamic relationship between BP and cerebral blood velocity (24, 26).

#### Middle cerebral artery blood velocity

MCAv was monitored with a 2-MHz pulsed transcranial Doppler ultrasound (Doppler Box; Compumedics DWL USA, Inc. San Juan Capistrano, CA). Identification and location of the left MCA was determined using standardized procedures (36). The probe was attached to a headset and secured with a custom-made headband and adhesive conductive ultrasonic gel (Tensive, Parker Laboratory, Fairfield, NY, USA) to ensure a stable position and angle of the probe throughout testing.

#### End-tidal partial pressure of carbon dioxide

End-tidal partial pressure of carbon dioxide (P_ET_CO_2_) were measured during the baseline period before the beginning of the exercise protocol (in men only) or the sit-to-stand (in women only) and squat-stand maneuvers (in both women and men) through a breath-by-breath gas analyzer (Breezesuite, MedGraphics Corp., MN) calibrated to known gas concentrations following manufacturer instructions before each evaluation.

#### Data acquisition

For each assessment, signals were analog-to-digital-converted at 1kHz via an analog-to-digital converter (Powerlab 16/30 ML880; ADInstruments, Colorado Springs, CO, USA) and stored for subsequent analysis using commercially available software (LabChart version 7.1; ADInstruments).

### Visit 1

#### Maximal oxygen consumption 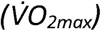

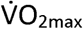 was determined during a progressive ramp exercise protocol performed on an electromagnetically braked upright cycle ergometer (Corival, Lode, the Netherlands). Following 3 min of rest, the evaluation started with 1 min of unloaded pedaling followed by an incremental ramp protocol (from 22 to 30 W/min according to participant’s history of training) to volitional exhaustion. Expired air was continuously recorded using a breath-by-breath gas analyzer (Breezesuite, MedGraphics Corp., MN, USA) for determination of 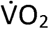, carbon dioxide production 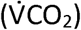, respiratory exchange ratio 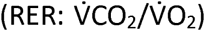 and P_ET_CO_2_. Maximal 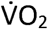 was defined as the highest 30-sec averaged 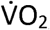, concurrent with a RER ≥ 1.15.

### Visit 2

#### Anthropometric measurements and resting hemodynamics

Stature and body mass were measured in each participant. Resting hemodynamic measurements included MAP (volume-clamp method using a finger cuff), which has been validated against intra-arterial pressure (19), heart rate (HR; ECG) and mean MCAv (MCAv_mean_) (transcraniaI Doppler ultrasound), which were continuously monitored on a beat-by-beat basis during 5 min of seated rest. Cerebrovascular conductance index (CVCi; MCAv_mean_/MAP) and its reciprocal, resistance (CVRi; MAP/MCAv_mean_) was then calculated. PetC0_2_ (breath-by-breath gaz analyzer) was continuously monitored (in women) on a breath-by-breath basis. The average values of the last 15 sec of recording represented the baseline. Since P_ET_CO_2_ was measured only in women during this 5 min of seated rest for technical reasons, the average values of the last 60 sec of P_ET_CO_2_ recording from the baseline period before the beginning of the exercise protocol (Visit 1) represented the baseline in men.

### Assessment of the dCA capacity and orthostatic tolerance

A multimetrics approach was employed to assess dCA to transient changes in MAP (Figure 1). We chose to force MAP oscillations using 2 separate techniques, to increase the input power (i.e. MAP) and improve the interpretation of the linear association between BP and MCAv (28).

**Figure 1.**
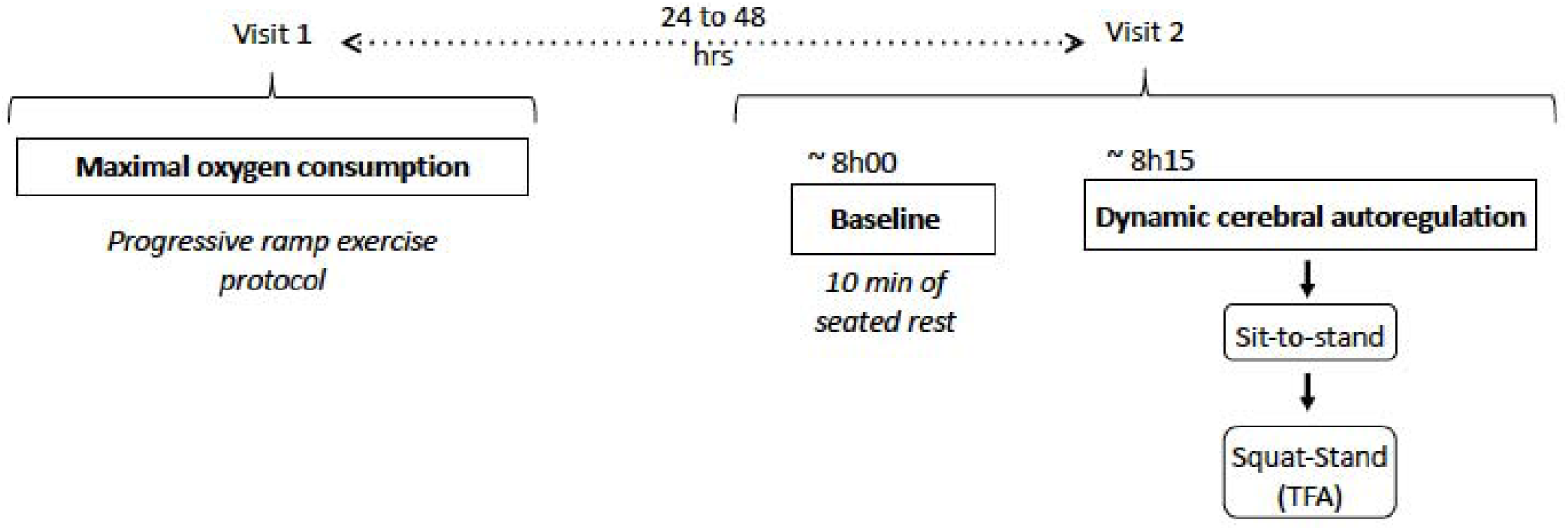
Experimental protocol

#### Sit-to-stand

Following 10 min of seated rest, participants rapidly (0-3s) stood up and maintained a standing position for 5 min without any movement or contraction of lower limb muscles. MAP and MCAv_mean_ were continuously monitored during this evaluation of the sit-to-stand response. P_ET_CO_2_ was measured only in women for technical reasons. In addition to the characterization of dCA, this technique permitted us to focus on the initial phase of the orthostatic response within the first 15 sec after standing. In addition, we examined the prevalence of IOH, defined as a decrease in systolic blood pressure ≥ 40 mmHg and/or a decrease in diastolic blood pressure ≥ 20 mmHg during the first 15 sec of standing (12).

#### Squat-stand maneuvers

For the squat-stand maneuvers, which were performed after a minimum of 10 min of rest, participants started in a standing position then squatted down until the back of their legs attained a ~ 90 degrees angle. This squat position was sustained for a specific time period, after which they moved to the standing position. Following instructions and practice, participants performed 5-min periods of repeated squat-stand maneuvers at a frequency of 0.05 Hz (10-sec squat, 10-sec standing) and 0.10 Hz (5-sec squat, 5-sec standing)(31). These large oscillations in MAP are extensively buffered by the cerebral vessels when executed at frequencies within the high-pass filter buffering range (<0.20 Hz) (38). The repeated squat-stand maneuver optimizes the signal-to-noise ratio enhancing the reproducibility and interpretability of findings through a physiologically-relevant MAP stimulus to the cerebrovasculature (28). The sequence of the squat-stand maneuvers was randomized between participants and each frequency was separated by 5 min of standing recovery. During these maneuvers, participants were instructed to maintain normal breathing and to avoid Valsalva. The linear aspect of the dynamic MAP-MCAv relationship was characterized via TFA (see the “Data analysis and statistical approach” section). MAP, HR, MCAv_mean_ and P_ET_CO_2_ were continuously monitored during this evaluation. An averaged P_ET_CO_2_ of the first and last five breaths of each maneuver (0.05 and 0.10 Hz) was calculated.

### dCA calculations

#### Acute cerebrovascular responses to hypotension induced by sit-to-stand

The following metrics were used to characterize the cerebral pressure-flow relationship to acute hypotension following the sit-to-stand,: 1) the reduction in MAP and MCAv_mean_ to their respective nadir (absolute: Δ MCAv_mean_, Δ MAP; and relative to baseline: Δ MCAv_mean_ (%), Δ MAP (%)); 2) the percent reduction in MCAv_mean_ per percent reduction in MAP (%ΔMCAv_mean_/%ΔMAP); 3) the time delay before the onset of the regulatory change; and 4) the rate of regulation (RoR).

The reduction in MAP and MCAv_mean_ is the difference between baseline MAP or MCAv_mean_ (averaged over the last 15 sec of seated rest before standing) and minimum MAP or MCAv_mean_ recorded after the sit-to-stand. %ΔMCAv_mean_/%ΔMAP upon standing was calculated as follows: [(baseline MCAv_mean_ - minimum MCAv_mean_)/(baseline MAP - minimum MAP)] × 100. The time delay before the onset of the regulatory change is the time lapse between the beginning of the sit-to-stand and the increase in CVCi (18). The onset of the regulatory response becomes visible when CVCi begins to continuously increase (without any subsequent transient reduction) following acute hypotension. This metric was assessed by two different observers (LL and PB). The physiological response to acute hypotension can be divided into two phases (21); Phase I is the time point after sit-to-stand where MCAv_mean_ changes are independent of any arterial baroreflex correction (1 to 7 s after sit-to-stand (13, 30, 33, 37). Phase II is the time point starting at the onset of arterial baroreflex and continuing for 4 sec (21). During Phase I, the rate of change in CVCi is directly related to dCA, without arterial baroreflex regulation (1). RoR was calculated during Phase I using the following equation:

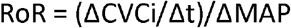

Where ΔCVCi/Δt is the linear regression slope between CVCi and time (t) during Phase I (a 2.5-s interval (Δt) after individually determined onset of the regulatory change following sit-to-stand was used for the analysis of RoR), and ΔMAP is calculated by subtracting baseline MAP from averaged MAP during Phase I (1, 21).

#### Assessment of the dynamic relationship between MAP and MCAv

Data was analyzed using commercially available software Ensemble (Version 1.0.0.14, Elucimed, Wellington, New Zealand) and are in accordance with the recommendations of the Cerebral Autoregulation Research Network (CARNet) (6). Beat-to-beat MAP and MCAv signals were spline interpolated and re-sampled at 4 Hz for spectral analysis and TFA based on the Welch algorithm. Each 5-min recording was first subdivided into 5 successive windows that overlapped by 50%. Data within each window were linearly detrended and passed through a Hanning window prior to discrete Fourier transform analysis. For TFA, the cross-spectrum between MAP and MCAv was determined and divided by the MAP auto-spectrum to derive the transfer function coherence (fraction of the MAP which is linearly related to MCAv), absolute gain (cm/sec/mmHg) (amplitude of MCAv change for a given oscillation in MAP), normalized gain (%/mmHg) and phase (radians) (difference of the timing of the MAP and MCAv waveforms).

The TFA coherence, gain and phase of the forced MAP oscillations were sampled at the point estimate of the driven frequency (0.05 and 0.10 Hz). These point estimates were selected as they are in the very low (0.02-0.07 Hz) and low (0.07-0.20 Hz) frequency ranges where dCA is thought to be most operant (28). Only the TFA phase and gain values where coherence exceeded 0.50 were included in analysis to ensure the measures were robust for subsequent analysis (29). Phase wrap-around was not present when coherence exceed 0.50 in the spontaneous data nor at any of the point-estimate values for squat-stand maneuvers.

#### Statistical analysis

Following the confirmation of normal distribution of data using Shapiro-Wilk W tests, between-group differences were analyzed using independent samples t-tests. Difference in the prevalence of IOH between groups was analyzed using the Fisher’s exact test. Relationships between variables were determined using Pearson product-moment. Statistical significance was established *a priori* at *p <0*.05 for all two-tailed tests. Data are expressed as mean ± standard deviation.

## RESULTS

### Dropout/Compliance

Three participants (two women and one men) were excluded from the sit-to-stand analysis because of an insufficient reduction in MAP (<10 mmHg) (29). The final sample size for the responses to acute hypotension following the sit-to-stand was 9 women and 10 men. Four participants were excluded from the TFA because of an inconsistent BP trace and a premature ending of the squat-stand maneuver related to the appearance of orthostatic symptoms (two women) or the absence of an appropriate ECG signal (two men). The final sample size for the TFA of forced oscillations in MAP and MCAv was 9 women and 9 men.

### Participant characteristics and baseline systemic and cerebrovascular hemodynamics

Age and BMI were comparable between groups. Volume of weekly training was also similar in both groups. Body mass, stature, P_ET_CO_2_ and 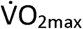 were lower in women (Table 1). Baseline MCAv_mean_ (+ 14cm sec^-1^; *p<0.001)* and CVCi (+ 0.14 cm·sec^-1^·mmHg^-1^; *p=0.002)* were higher, whereas CVRi was lower (−0.29 mmHg· cm·sec^-1^ *p=0.001),* in women. All other baseline systemic hemodynamics were similar between groups (Table 1).

**Table 1.** Baseline characteristics and resting values between women and men

| Baseline characteristics | Women | Men | p values |
| --- | --- | --- | --- |
| n | 11 | 11 |  |
| Age (y) | 25 ± 4 | 24 ± 2 | 0.610 |
| Stature (m) | 1.64 ± 0.07 | 1.78 ± 0.09 | <0.001 |
| Body mass (kg) | 61.0 ± 5.7 | 72.5 ± 10.4 | 0.007 |
| Body mass index (kg/m<sup>2</sup>) | 23 ± 2 | 23 ± 2 | 0.992 |
| Training volume (min) | 466 ± 151 | 511 ± 188 | 0.548 |
| Maximal oxygen uptake (ml/kg·min<sup>-1</sup>) | 48.1 ± 4.1 | 56.8 ± 4.4 | <0.001 |
| **Resting values** |  |  |  |
| n | 11 | 11 |  |
| Heart rate (bpm) | 73 ± 11 | 74 ± 12<br>(n=10) | 0.853 |
| Mean arterial pressure (mmHg) | 96 ± 10 | 97 ± 9 | 0.919 |
| Middle cerebral artery mean blood velocity (cm sec<sup>-1</sup>) | 75 ± 7 | 61 ± 7 | <0.001 |
| Cerebrovascular resistance index (mmHg·cm·sec<sup>-1</sup>) | 1.30 ± 0.16 | 1.59 ± 0.20 | 0.001 |
| Cerebrovascular conductance index (cm·sec<sup>-1</sup>·mmHg<sup>-1</sup>) | 0.78 ± 0.10 | 0.64 ± 0.08 | 0.002 |
| End-tidal partial pressure of carbon dioxide (mmHg) | 36.9 ± 2.3 | 42.5 ± 3.5 | <0.001 |

Data are presented as mean ± SD.

### Influence of sex on dCA

#### Responses to acute hypotension following the sit-to-stand

MAP (96 ± 10 vs. 97 ± 9 mmHg; *p= 0.919)* was comparable between groups and MCAv_mean_ (75 ± 7 vs. 61 ± 7 cm sec^-1^; *p=0.0001)* was higher in women than men at baseline. Upon standing, although the reduction [absolute (Figure 3) and relative (−26 ± 4 vs. -25 ± 11 %; *p=0.862*)] in MAP to nadir was similar between women and men, the absolute decrease in MCAv_mean_ was of greater amplitude (−20 ± 8 vs. -11 ± 7 cm sec^-1^; *p=0.018).* There were no group differences in the time taken to reach the nadir for MAP (7 ± 1 vs. 7 ± 2 sec; p=0.510). The time delay before the onset of the regulatory change was almost two-fold longer in women *(p=0.007;* Figure 3) following the sit-to-stand. After the onset of the regulatory response, RoR was not different between groups (0.26 ± 0.27 vs. 0.23 ± 0.36 sec^-1^; p=*0.867*). In women, mean change in P_ET_CO_2_ from baseline to average MAP during the first 15 sec following the sit-to-stand was -0.8 ± 1.1 mmHg.

**Figure 2.**
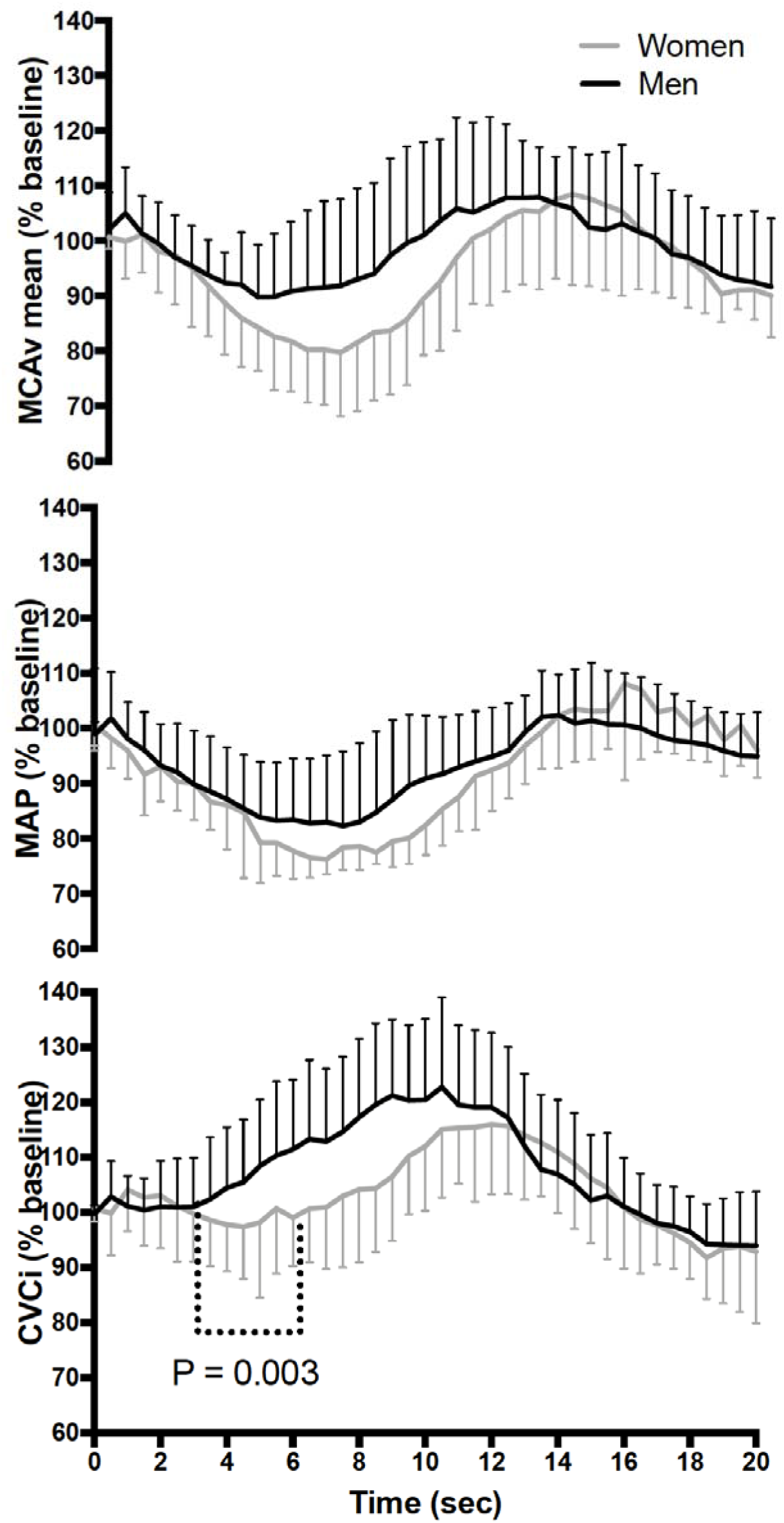
Normalized response of mean arterial pressure (MAP), middle cerebral artery mean blood velocity (MCAv_mean_) and cerebrovascular conductance index (CVCi) in women (gray line) and men (black line) following sit-to-stand. Time 0 indicates the transition from sitting to standing.

**Figure 3.**
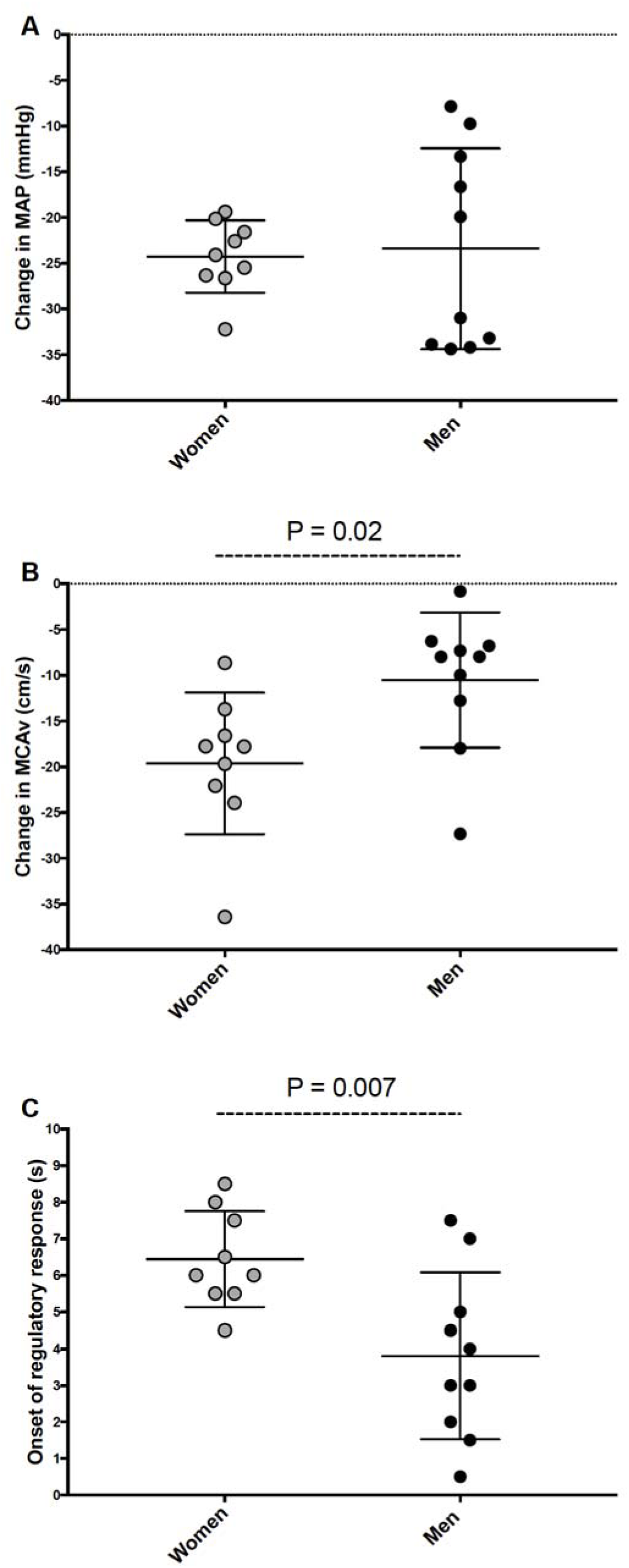
Cerebrovascular responses following sit-to-stand. Change in middle cerebral artery mean blood velocity (MCAv_mean_) from baseline (A), relative change in MCAv_mean_ on the relative change in MAP (B), onset of the regulatory response (C) and rate of regulation (D). Shaded circles indicate women and black circles, men.

#### TFA of forced oscillations in MAP and MCAv

MAP and MCAv power spectrum densities (0.05 and 0.10 Hz) during forced oscillations were not different between women and men (Table 2). Coherence during 0.05 Hz squat-stand was lower in women than men (*p=0.038*). Absolute TFA gain during 0.05 Hz squat-stand was higher in women compared to men (*p=0.017*). All the other metrics were not different between women and men (Table 2 and Figure 4). Changes in P_ET_CO_2_ from the beginning of each squat-stand maneuver (0.05 Hz: +1.6 ± 1.2 vs. +1.4 ± 2.3 mmHg; *p=0.50* and 0.10 Hz: +1.3 ± 0.7 vs. +2.0 ± 3.2 mmHg *p=0.66*) were comparable between women and men.

**Table 2.**
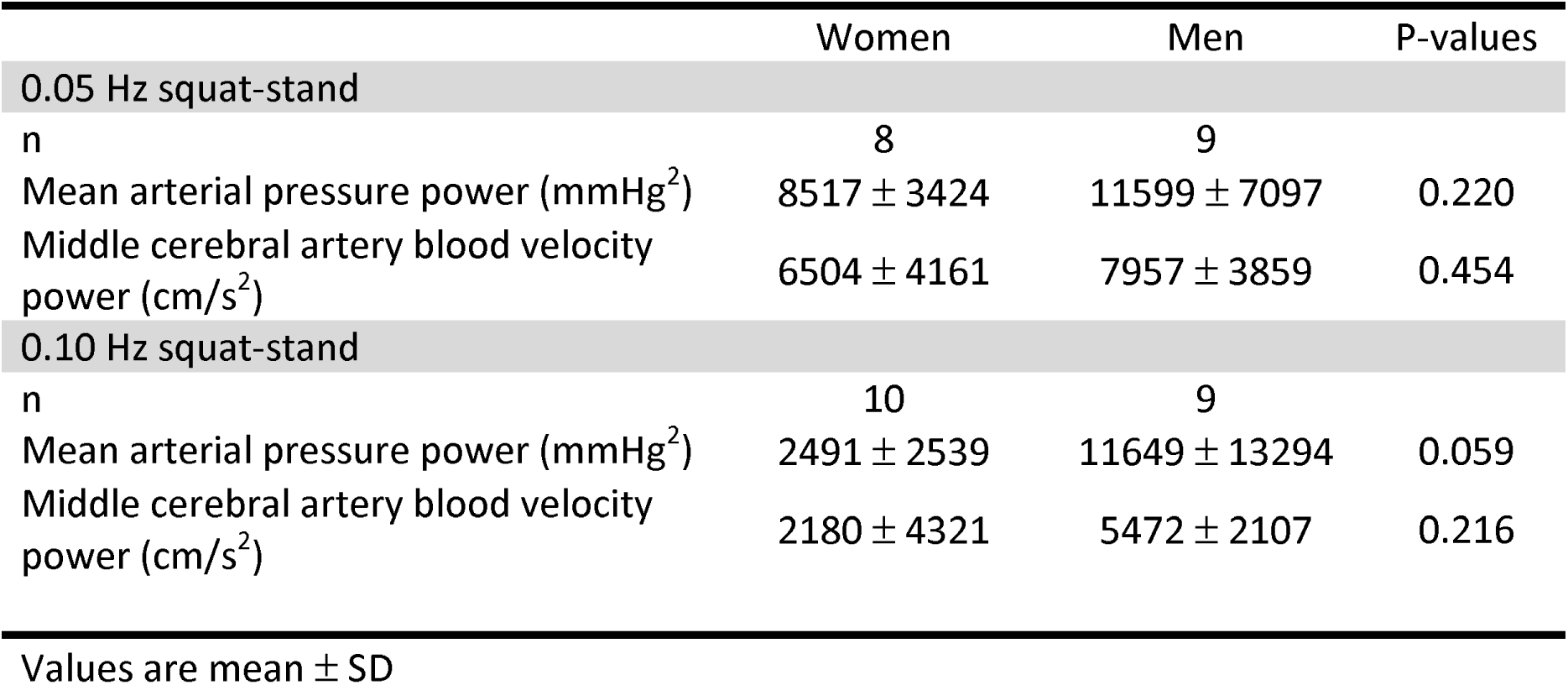
Power spectrum densities of forced oscillations in mean arterial pressure and middle cerebral artery blood velocity during squat-stand maneuvers

|  | Women | Men | P-values |
| --- | --- | --- | --- |
| **0.05 Hz squat-stand** |  |  |  |
| n | 8 | 9 |  |
| Mean arterial pressure power (mmHg<sup>2</sup>) | 8517 ± 3424 | 11599 ± 7097 | 0.220 |
| Middle cerebral artery blood velocity power (cm/s<sup>2</sup>) | 6504 ± 4161 | 7957 ± 3859 | 0.454 |
| **0.10 Hz squat-stand** |  |  |  |
| n | 10 | 9 |  |
| Mean arterial pressure power (mmHg<sup>2</sup>) | 2491 ± 2539 | 11649 ± 13294 | 0.059 |
| Middle cerebral artery blood velocity power (cm/s<sup>2</sup>) | 2180 ± 4321 | 5472 ± 2107 | 0.216 |

Values are mean ± SD

**Figure 4.**
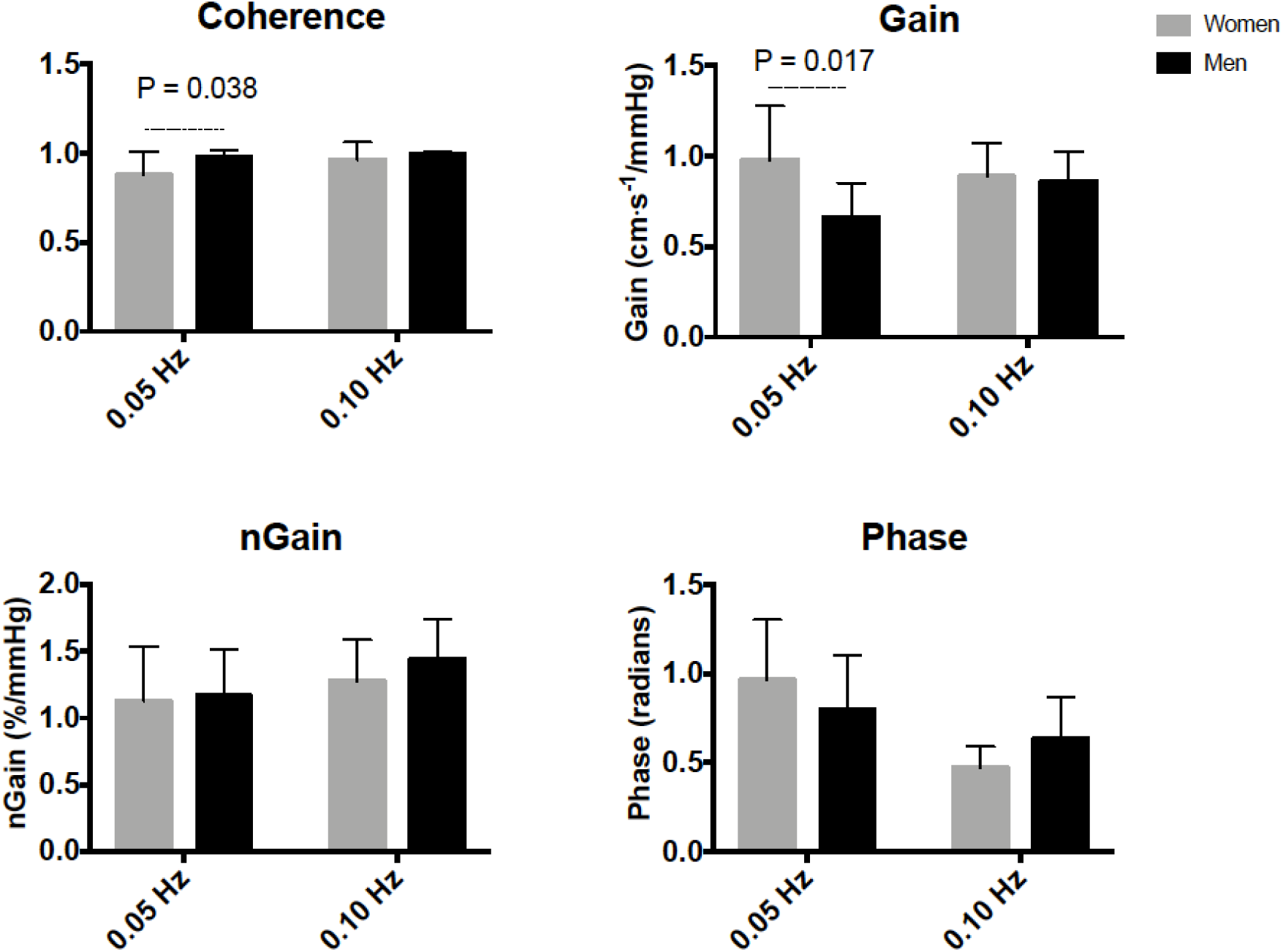
Transfer function analysis of forced oscillation in mean arterial pressure and middle cerebral artery blood velocity. Group averaged coherence, phase, gain and normalized gain (nGain) for 0.05 Hz and 0.10 Hz squat-stand.

### Initial orthostatic hypotension

The prevalence of IOH (women: 4/9 vs. men: 5/9, *p=0.637*) was not different between groups and neurogenic syncope-related symptoms were not reported by the participants who experienced IOH. There were no correlations between metrics of dCA and decreases in MAP or MCAv_mean_ to their nadir upon standing (data not shown).

## DISCUSSION

The main findings of this study were three-fold: healthy active women had (1) a delayed onset of their cerebral autoregulatory response; (2) a greater decrease in MCAv_mean_ in response to transient hypotension induced by a sit-to-stand; and (3) higher absolute TFA gain during 0.05 Hz repeated squat-stand maneuvers. Taken together, these findings imply that the brain vasculature of healthy women has a reduced ability to dampen fast and large MAP oscillations compared to men. Finally, the prevalence of orthostatic intolerance was similar in women and men, and there were no associations between dCA and orthostatic tolerance metrics in our women. Overall, these findings support the notion that despite dCA changes in women, they appear to be functionally inert, in that IOH was not different between groups.

### Resting cerebral hemodynamics

In healthy young adults, resting CBF is higher in women compared to age-matched men (11, 20, 21, 30). It has been speculated this CBF, when monitored using transcranial (intra-cranial arteries) or duplex Doppler (extra-cranial arteries) ultrasound, is associated with circulating ovarian hormones and the menstrual cycle (4, 16, 17). The current results of greater resting MCAv_mean_ and CVCi in women are in agreement with this notion (Table 1).

### dCA

#### Acute hypotension induced by sit-to-stand

The literature related to the influence of sex on the cerebrovascular response to BP changes is sparse and limited to just a few variables. To the best of the authors knowledge currently only one study has shown differential regulation between women and men in response to a sit-to-stand maneuver (9). Deegan et al. showed that women had improved dCA metrics with higher autoregulation index (ARI), smaller reductions in MAP and MCAv_mean_ as well as a lower reduction in %ΔMCAv_mean_/ %ΔMAP. However, subjects were aged over 70 years old, so the potential confounding effects of circulating sex hormones could be discounted (9). In younger individuals, men seem to possess a better static CA during head-up tilt (35) and higher cerebrovascular resistance while standing than women (2). Conversely, others reported no sexual differences in CBF regulation during head-up tilt (35) or the sit-to-stand maneuver (11).

In the present study, we demonstrate an impairment in the cerebrovasculature of healthy women when this system is challenged via a large and rapid reductions in BP. Despite a comparable MAP reduction between groups, women had a delayed onset of the regulatory response, as well as a larger reduction in MCAv_mean_ following sit-to-stand (Figure 2). Differences in analytical techniques might explain disparities between the current results and those from the broader literature. In fact, in two studies where a sit-to-stand was included (2, 14) the sitting and standing positions were compared during the steady-state phases (in contrast to the current design which assessed the dynamic phase) associated with the posture change. By comparing only the averaged data of the last minute at each position in their healthy men and women (5 min seated and 10 min standing) (2), variables (i.e. HR, MAP, MCAv_mean_, cerebrovascular resistance) will have had sufficient time for the baroreceptors to have adjusted and enabled recovery from the acute hypotension, thus reducing the acute influence of the orthostatic challenge on the metrics of interest. The discrepancy between the current findings and the previous literature further emphasizes the importance of analysing the dynamic response of the sit-to-stand (i.e. dCA) instead of simply comparing steady-state hemodynamics before and after a given BP challenge (i.e. static CA). Therefore, it is important to consider the influence of various measures during both static and dynamic cerebral autoregulatory challenges (31) (such as %ΔMCAv_mean_ /%ΔMAP, the onset of the regulatory response and RoR) when assessing sexual differences in the CBF response to a sit-to-stand. Also, changes in P_ET_CO_2_ during the first 15 sec upon standing was minimal in women in the current study. Although this variable could not be measured in men for technical reasons, sex does not seem to influence the change in P_ET_CO_2_ during a sit-to-stand, at least in an older population (8).

#### TFA of spontaneous and forced oscillations

The literature examining the influence of sex on TFA of spontaneous and forced oscillations is limited. To the best of the authors knowledge, only three studies have used TFA to examine sex differences in dCA. Using spontaneous oscillations with participants in the seated position, Deegan et al. (9) reported reduced HF gain in women compared to men, suggesting an enhanced dCA in women. However, participants from this study were considerably older (> 70 years old) than the current investigation. In a population ranging in age from 21 to 80 years old, Xing et al. (37) reported lower VLF gain and higher phase during 0.05 Hz repeated squat-stand maneuvers as compared with men, again suggesting a better dCA in women. In contrast to these two other investigations, Patel *et al.* found no differences in dCA between sexes using TFA of spontaneous oscillations in 129 participants with a mean age of ∼57 years (25).

The current study revealed women had a higher gain during 0.05 Hz repeated squats-stand maneuvers (Figure 4) indicative of a more compliant cerebrovasculature compared to men. The higher gain during 0.05 Hz repeated squats-stand maneuvers also relates to the lower MAP power spectrum densities at 0.05 Hz as an increased gain means there is a greater change in MCAv for a given change in MAP. Accordingly, the lower MAP power spectrum densities in women with a similar MCAv power spectrum densities between sexes during 0.05 Hz squat-stand would indicate more of the BP being transferred to the women brain. Discrepancies between the current results and the previous literature might come from the age differences between our studied populations. Many of the women included in the previous studies were probably post-menopausal, which could attenuate the influence of hormone differences in dCA between sexes (9). Consistent with this notion are previous reports which demonstrated there were no sexual differences in cerebral hemodynamics between old women and young men (11, 20). Further research is needed to better understand the influence of the menstrual cycle on TFA metrics. Of note, P_ET_CO_2_ is unlikely to explain the reported differences in dCA metrics gathered during forced oscillations in BP considering that small changes in P_ET_CO_2_ during repeated squat-stand maneuvers were not different between women and men.

### To what extent are the differences in dCA in healthy active women physiologically / clinically meaningful?

In the current study, the prevalence of IOH was similar between women and men, even though women showed a diminished dCA and a greater reduction in MCAv_mean_ upon standing (Figures 3 and 4). However, the current results did not correlate dCA metrics and reductions in MAP and MCAv_mean_ during orthostatic stress induced by the sit-to-stand in in women. These results suggest the subtle changes in dCA in healthy young women do not translate into functional outcome, at least in response to a sit-to-stand. Our findings thus bring into question the physiological validity of the subtle dCA differences which have been observed in otherwise healthy young women. These dCA changes may simply not be important enough to influence orthostatic tolerance, as the alterations in both the sit-to-stand and TFA data indicate the cerebrovasculature of the women may be more compliant and be able to better withstand alterations in BP induced in the current investigation. Larger reductions in MAP may thus be necessary to affect orthostatic tolerance in the presence of functional impairments in dCA. Considering the higher resting CBF reported in the current study and by others (2, 11, 21), the absence of associations between the attenuated dCA and the prevalence of IOH reported here could be related to a higher cerebrovascular reserve in women to face a rapid transient hypotension induced by a sit-to-stand compared to men. These dCA changes could then become clinically meaningful with a larger BP reduction.

### Limitations

Some limitations to our study deserve further discussion. Only young healthy and fit women and men participated to this study and the results cannot be generalized to other populations (such as older individuals or hypertensive patients). Considering the presence of hysteresis in the cerebral pressure-flow relationship (5, 23, 33), these results cannot be translated to situations involving transient hypertension.

P_ET_CO_2_ was not available in men during the sit-to-stand. Baseline P_ET_CO_2_ in men was thus determined from the baseline period before the beginning of the exercise protocol. Although this could have partly explained the difference in baseline P_ET_CO_2_ between groups, women had comparable P_ET_CO_2_ values during the baseline period before the beginning of the exercise protocol (Visit 1) and the 5 min of baseline rest (Visit 2) (36.9 ± 2.3 vs. 36.5 ± 2.8 mmHg *p=0.47*). Men most likely had similar P_ET_CO_2_ values between visits as well. Of note, such a difference in resting P_ET_CO_2_ between women and men has already been reported (10). In addition, changes in P_ET_CO_2_ during this maneuver was minimal in women and sex does not seem to influence the change in P_ET_CO_2_ during a sit-to-stand in an older population (8). This variable most likely played a minimal role in the metrics related to the sit-to-stand. Indeed, since women had lower P_ET_CO_2_ at baseline (which would lead to vasoconstriction), it did not play a role in the augmented gain values for either the sit-to-stand nor TFA parameters. A reduction in CO_2_ should cause a decrease in gain and this was not observed. Instead, the altered CO_2_ at baseline likely did not cause the noted differences in the study and in fact may have instead limited the extent of them.

Further to this point, cerebral blood velocity in the MCA was measured with transcranial Doppler ultrasound and is representative of flow if the diameter of the arteries remains constant. Changes in MAP and P_ET_CO_2_ have been associated with changes in the diameter of the internal carotid artery and MCA. However, the physiological range of variation in MAP and P_ET_CO_2_ from the current study will most likely be associated with a minor effect on the diameter of the MCA (19, 34).

Considering the women in the current investigation included: a group of women taking oral contraceptives continuously (n=2), having an intrauterine device (n=2) or were assessed during days 1-10 of their menstrual cycle (n=7), we are unable to ascertain if the elevated CBF in the current investigation was influenced by the oscillatory nature of these hormones throughout the menstrual cycle. Further research is warranted to determine the specific effects the stages of the menstrual cycle play on these measures. In addition, we cannot rule out the possibility that dCA changes reported in the current study could be amplified and ultimately affect orthostatic tolerance later in the menstrual cycle. Indeed, pre-syncopal symptoms are more frequent during the menses phase of the menstrual cycle (22). Menses usually last from days 0 to 5. However, some of our women were tested after day 5. Therefore, their orthostatic tolerance could be better following the menses phase leading to a lower prevalence of orthostatic hypotension and less symptoms of cerebral hypoperfusion in our cohort. Of note, Abidi et al. recently assessed static CA and peripheral hemodynamics during a sit-to-stand protocol in oral contraceptives and non-oral contraceptives users, as well as during the high and low hormones phases. They reported differences in MAP regulation, but no impact of the menstrual cycle or oral contraceptives use on the cerebrovascular response (2).

Although comparable in terms of previous weekly training volume, women and men were not matched for 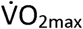 and it could have influenced our results. We have recently reported a reduced dCA with elevated CRF when the brain is challenged with large and rapid MAP change in men (18). We speculate the differences in dCA metrics observed between women and men in the current study would be even more important if women had a higher CRF to match men’s level. However, since CRF is usually superior in men vs. women for a similar charge of training, a matching of 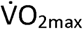 between male and female athletes would not represent a real-life situation (7). Either women would be too trained or men untrained.

## CONCLUSION

These results indicate the cerebrovasculature of healthy fit women has an attenuated ability to react to changes in BP compared to men, when the brain is challenged with large and rapid BP oscillations. However, these subtle dCA changes are not translated into functional impairments using initial orthostatic hypotension as the functional outcome, which could be related to a higher cerebrovascular reserve in women to face a rapid transient hypotension induced by a sit-to-stand compared to men.

## GRANTS

This study has been supported by the Ministère de l’Éducation, du Loisir et du Sport du Québec and the Foundation of the Institut universitaire de cardiologie et de pneumologie de Québec-Université Laval. L.L. is supported by a doctoral training scholarship from the Société québécoise d’hypertension artérielle. S.I. is supported by a doctoral training scholarship from the Fonds de recherche du Québec-Santé (FRQS). D.M.B. is a Royal Society Wolfson Research Fellow (#WM170007).

Authors contribution
P.B. contributed to the original idea of the study; L.L.,K.R, S.I, M.P.,O.L., S.M. and D.A. contributed to data collection; L.L. contributed to data analyses; L.L., J.D.S., D.M.B. and P.B. contributed to data interpretation; L.L.,J.D.S, D.M.B and P.B drafted the article. All authors provided approval of the final article.

